# BiPOLES: a tool for bidirectional dual-color optogenetic control of neurons

**DOI:** 10.1101/2020.07.15.204347

**Authors:** Johannes Vierock, Silvia Rodriguez-Rozada, Florian Pieper, Alexander Dieter, Amelie Bergs, Nadja Zeitzschel, Joachim Ahlbeck, Kathrin Sauter, Alexander Gottschalk, Andreas K. Engel, Peter Hegemann, J. Simon Wiegert

## Abstract

Optogenetic manipulation of neuronal activity has become an indispensable experimental strategy in neuroscience research. A large repertoire of excitatory and inhibitory tools allows precise activation or inhibition of genetically targetable neuronal populations. However, an optogenetic tool for reliable bidirectional control of neuronal activity allowing both up- and downregulation of the same neurons in a single experiment is still missing. Here we report BiPOLES, an optogenetic tool for potent excitation and inhibition of the same population of neurons with light of two different colors. BiPOLES consists of an inhibitory, blue-light-sensitive anion-conducting channelrhodopsin fused to an excitatory, red-light-sensitive cation-conducting channelrhodopsin in a single, trafficking-optimized tandem protein. BiPOLES enables multiple new applications including potent dual-color spiking and silencing of the same neurons *in vivo* and dual-color optogenetic control of two independent neuronal populations.

## Introduction

To prove necessity and sufficiency of a particular neuronal population for a specific behavior, a cognitive task, or a pathological condition, it is desirable to both faithfully inhibit and activate this exact same population of neurons. In principle, optogenetic manipulations should allow such interventions. However, excitation and inhibition of the neuronal population of interest is commonly done in separate experiments, expressing either an excitatory or an inhibitory opsin. Alternatively, if both opsins are co-expressed in the same cells, it is essential to achieve efficient membrane trafficking of the two opsins, equal subcellular distribution and a defined ratio between excitatory and inhibitory action at the respective wavelengths, so that neuronal activation and silencing can be precisely controlled in all transduced cells. This is particularly challenging when AAV-transduction of the optogenetic actuators is required *in vivo*.

A second, long-standing challenge is the independent activation of two defined neuronal populations. Although two spectrally distinct opsins have been combined previously to spike two distinct sets of neurons^1-4^, careful calibration and dosing of blue light is required to avoid activation of the red-shifted opsin, as all rhodopsins are activated to a certain extent by blue light. This typically leaves only a narrow spectral and energetic window to activate the blue-but not the red-light-sensitive opsin. Thus, dual-color control of neurons is particularly challenging in the mammalian brain where light intensities decrease by orders of magnitude over a few millimeters in a wavelength-dependent manner^5,6^.

Capitalizing on the recent advent of anion-conducting channelrhodopsins (ACRs)^7-9^ and a previously established tandem gene-fusion strategy^10^, we generated BiPOLES, a <u>Bi</u>directional <u>P</u>air of <u>O</u>psins for <u>L</u>ight-induced <u>E</u>xcitation and <u>S</u>ilencing. First, BiPOLES enables potent, light-mediated silencing and activation of the same neurons *in vivo* by a single optogenetic tool and second, dual-color control of two distinct neuronal populations without cross-talk at light intensities spanning multiple orders of magnitude, when combined with a second blue-light-sensitive channelrhodopsin (ChR).

## Results

As a general strategy, we fused the red-light-activated cation channel Chrimson with various blue-or green-light-activated ACRs. This approach aimed for colocalized and balanced hyper-and depolarization with blue and red light starting from a physiological membrane voltage as well as restriction of the depolarizing light spectrum to a narrow, orange-red window as the ACR compensates the blue-light-activated Chrimson currents. The opsins were linked by sequences composed of the Kir2.1 membrane trafficking signal (ts)^11^, a cyan or yellow fluorescent protein, and the transmembrane β helix of the rat gastric H+/K+ ATPase (βHK) to maintain correct membrane topology of both opsins^10^ (Fig. 1A).

**Figure 1:**
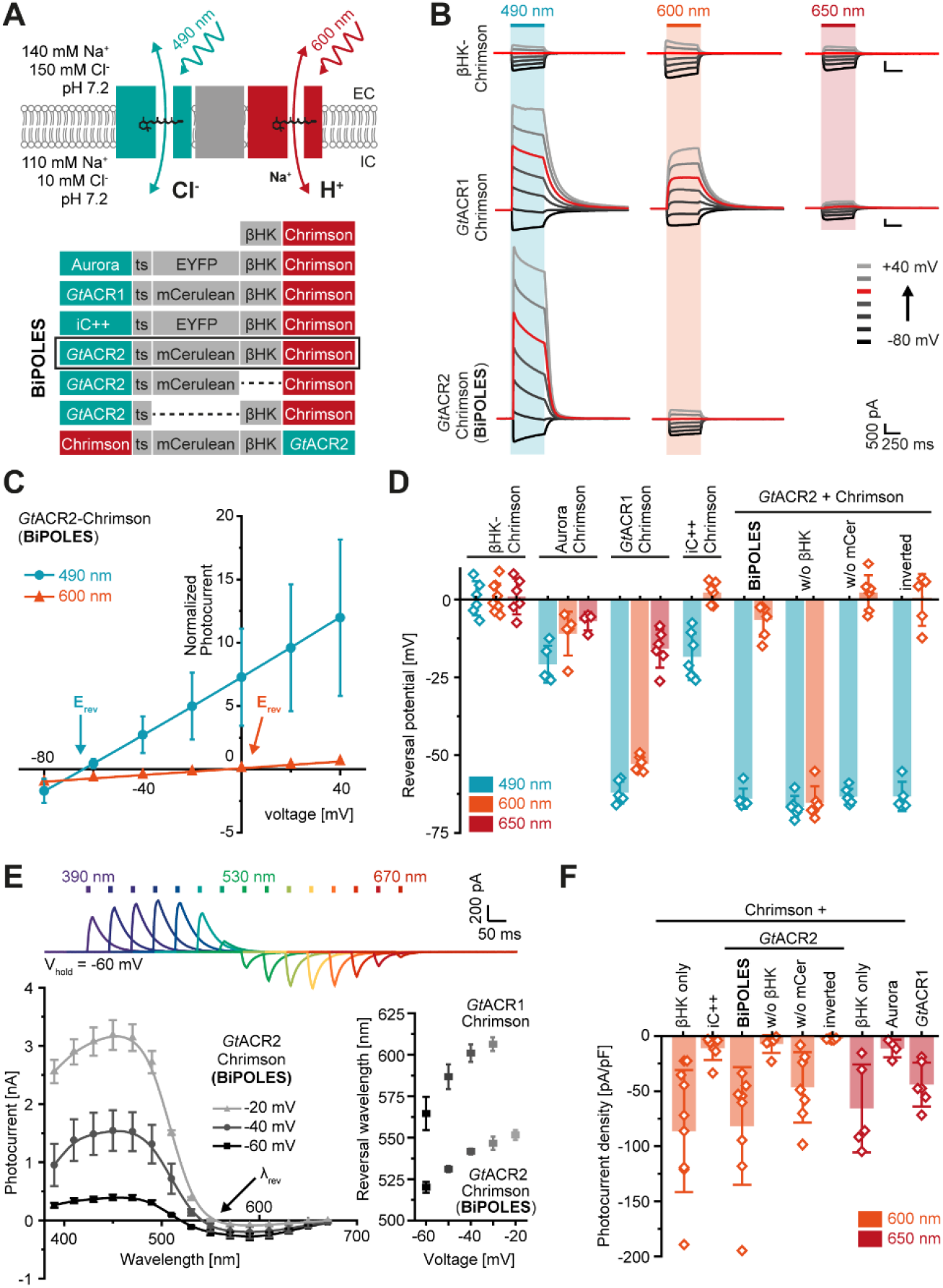
Development of BiPOLES and biophysical characterization. **(A)** Molecular scheme of BiPOLES with the extracellular (EC) and intracellular (IC) ionic conditions used for HEK293-cell recordings. The blue-green-light-activated natural anion channels *Gt*ACR1 and *Gt*ACR2 or the engineered ChR-chimeras iC++ and Aurora were fused to the red-light-activated cation-conducting Chrimson by a transmembrane spanning linker region consisting of a trafficking signal (ts), a yellow or cyan fluorescent protein (EYFP, mCerulean3) and the βHK transmembrane fragment. The fusion construct termed BiPOLES is indicated by a black frame. **(B)** Representative photocurrents of βHK-Chrimson-mCerulean (top), *Gt*ACR1-ts-mCerulean-βHK-Chrimson (middle) *Gt*ACR2-ts-mCerulean-βHK-Chrimson (BiPOLES, bottom) in whole-cell patch clamp recordings from HEK293 cells at 490, 600 and 650 nm illumination. **(C)** Normalized peak photocurrents of BiPOLES at different membrane voltages evoked at either 490 or 600 nm (see panel B, mean ± SD; n = 6 - 8; normalized to the peak photocurrent at −80 mV and 600 nm illumination). **(D)** Reversal potential of early peak photocurrents during 500-ms illumination with 490, 600, or 650 nm light as shown in (B) (mean ± SD; n = 5 - 8). **(E)** Top: Representative photocurrents of BiPOLES with 10 ms light pulses of different color and equal photon flux at −60 mV. Lower left: Action spectra of BiPOLES at different membrane voltages (λ_rev_ = photocurrent reversal wavelength, mean ± SEM, n = 4 - 9). Lower right: λ_rev_ of *Gt*ACR1-ts-mCerulean-βHK-Chrimson and BiPOLES at different membrane voltages (mean ± SD; n = 3 - 9). **(F)** Peak photocurrent densities at −80 mV and 600 nm or 650 nm illumination as shown in (B) (Mean ± SD; n = 5 - 8).

All ACR-Chrimson tandems were evaluated in human embryonic kidney cells (HEK) under matched experimental conditions. In all constructs, except the one lacking the βHK-subunit, blue-light-activated currents shifted towards the chloride Nernst potential whereas red-light-activated currents shifted towards the proton Nernst potential (Fig.1B-D, S1). Reversal potentials varied strongly for the different tandem variants indicating considerable differences in the wavelength-specific anion/cation conductance ratio (Fig.1D). At a defined membrane potential between the Nernst potential for chloride or protons, blue and red light induced outward and inward currents, respectively. The specific wavelength of photocurrent inversion (λ_rev_) depended on the action spectrum of the ACR, the relative conductance of the ACR and Chrimson, and the relative driving force for anions, cations and protons defined by the membrane voltage and the respective ion gradients (Fig.1E).

*Gt*ACR2-ts-mCerulean-βHK-Chrimson – from here on termed BiPOLES – was the most promising variant, showing first, the largest difference in reversal potential upon blue-or red-light excitation (Fig.1D), second, equal inward and outward currents at −60 mV, which is near the resting membrane voltage (Fig.1E) and, third, the highest red-light-activated photocurrents, comparable to those of Chrimson expressed alone (Fig.1F).

Next, we validated the bidirectional action and the applicability of BiPOLES as an optogenetic tool in CA1 pyramidal neurons of rat hippocampal slice cultures. We observed membrane-localized BiPOLES expression, most strongly in the somatodendritic compartment (Fig. 2A). Illumination triggered large photocurrents with biophysical properties similar to those observed in HEK cells (Fig. S2). To enhance membrane trafficking and to avoid axonal localization of BiPOLES, we generated a soma-targeted variant (somBiPOLES) by attaching a C-terminal Kv2.1-trafficking sequence^12^. somBiPOLES showed improved membrane localization (Fig. 2A, S9A), enhanced photocurrents (Fig. S3) and higher light sensitivity compared to BiPOLES (Fig. S4). In current-clamp recordings orange and red light reliably induced action potentials (APs) in somBiPOLES expressing cells with a similar efficacy as Chrimson (Fig. 2B, S4). Notably, blue light up to 100 mW/mm^2^ did not trigger APs due to robust shunting by *Gt*ACR2, whereas Chrimson alone induced spikes also with blue light above 0.95 mW/mm^2^ (Fig. 2B, S4). Next, we tested the silencing capabilities of BiPOLES and somBiPOLES by measuring their ability to shift the rheobase (see Methods) and suppress APs with blue light. Both variants significantly shifted the rheobase towards larger currents at intensities above 0.1 mW/mm^2^ (Fig. 2C, S5), with somBiPOLES showing complete spike block in some cases. We further demonstrate the potent silencing capacity of somBiPOLES and BiPOLES by combining blue-and red-light pulses showing that red-light-evoked spikes were reliably inhibited with a coinciding blue-light pulse (Fig. 2D). Moreover, simultaneous illumination with blue and orange light at varying ratios enabled dynamic clamping of the neuronal membrane voltage (Fig. S6). Taken together, BiPOLES and somBiPOLES enable efficient, bidirectional control of neuronal activity.

**Figure 2:**
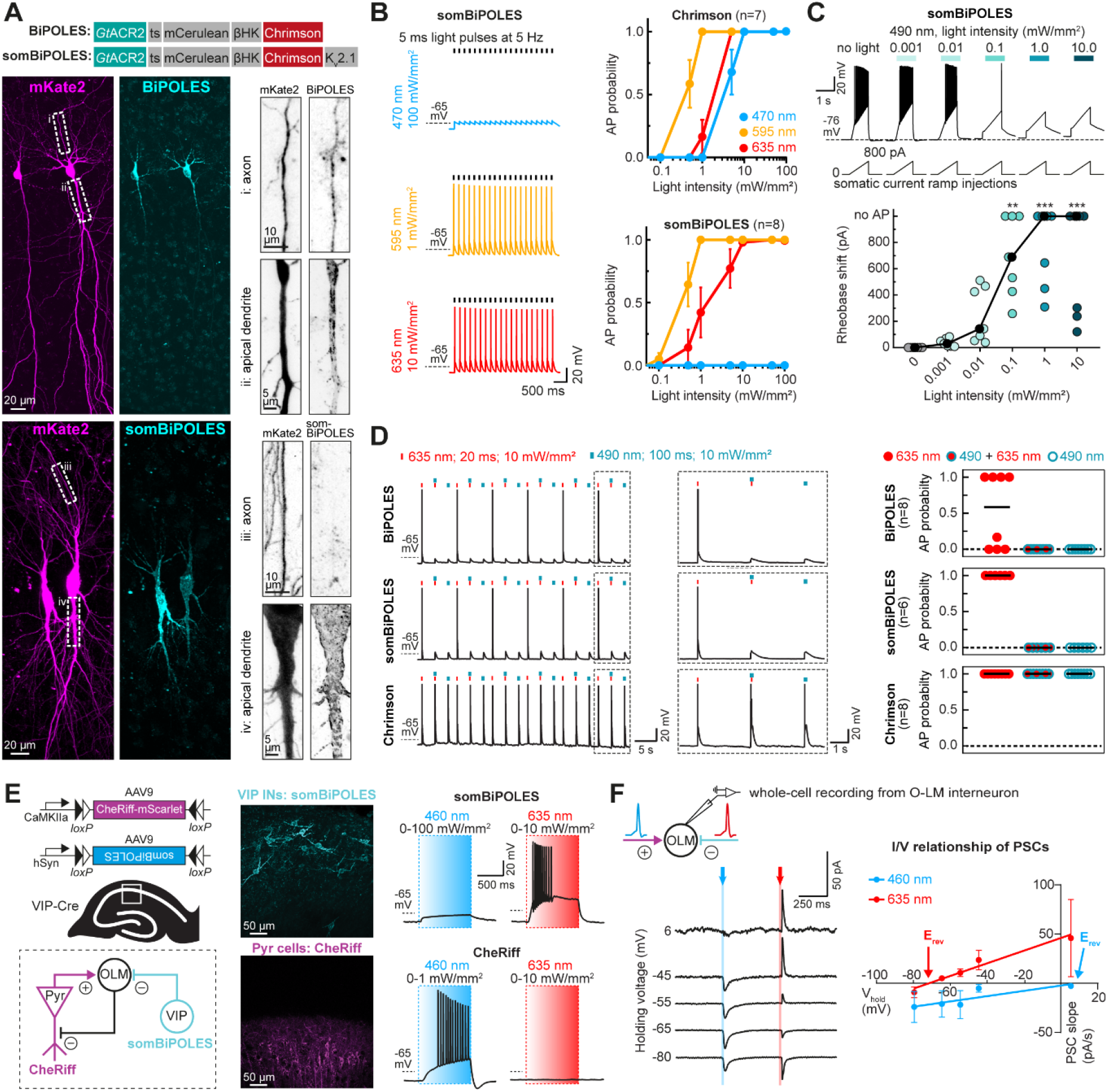
Excitation and silencing of hippocampal neurons with BiPOLES and somBiPOLES. **(A)** Molecular scheme of BiPOLES and somBiPOLES as used in neurons and maximum-intensity projection images of 2-photon stacks showing expression of BiPOLES (top) or soma-targeted BiPOLES (somBiPOLES, bottom) co-expressed with mKate2 in CA1 or CA3 pyramidal neurons of organotypic hippocampal slices. Magnified views of axonal or somato-dendritic compartments are shown as inverted gray-scale images. Note absence of somBiPOLES in the axon. **(B)** Current-clamp (IC) characterization of somBiPOLES and Chrimson in CA1 pyramidal cells to determine light-evoked action potential (AP)-probability at different wavelengths. Left: Example traces. Right: quantification of light-mediated AP probability at indicated wavelengths and intensities (mean ± SEM, n = 7 - 8). **(C)** IC characterization of somBiPOLES-mediated neuronal silencing. Current ramps (from 0–100 to 0–900 pA) were injected into somBiPOLES-expressing CA1 pyramidal cells to induce APs during illumination with blue light at indicated intensities (from 0.001 to 10 mW/mm^2^, black circles: medians, n = 7, Friedman test, **p < 0.01, ***p < 0.001). **(D)** IC characterization of bidirectional optical spiking-control with BiPOLES and somBiPOLES. Left: Voltage traces showing red-light-evoked APs, which were blocked by a concomitant blue light pulse in (som)BiPOLES expressing cells. Right: quantification of AP probability under indicated conditions (black horizontal lines: medians, n = 6 - 8). **(E)** Independent dual-color control of two neuronal populations with somBiPOLES and CheRiff. Left: strategy to achieve mutually exclusive expression of CheRiff-mScarlet in CA1 pyramidal neurons and somBiPOLES in VIP-positive GABAergic neurons. Both cell types innervate O-LM interneurons in CA1. Middle: Maximum-intensity projection images of 2-photon stacks showing expression of somBiPOLES in VIP-interneurons (cyan) and CheRiff-mScarlet in the pyramidal layer of CA1 (magenta). Right: IC-recordings demonstrating mutually exclusive spiking of somBiPOLES-and CheRiff-expressing neurons under red or blue illumination (see fig. S4 for details). **(F)** Postsynaptic whole-cell voltage-clamp recordings at indicated membrane voltages showing EPSCs and IPSCs upon blue-and red-light pulses, respectively. Right: quantification of blue-and red-light-evoked PSCs and their reversal potential (mean ± SEM, n = 7 - 8).

Since BiPOLES permits neuronal spiking exclusively with orange-red light, this opens new possibilities for two-color excitation of genetically distinct but spatially intermingled neuronal populations using a second, blue-light-activated ChR. To demonstrate this, we expressed somBiPOLES in CA1 VIP interneurons and CheRiff, a blue-sensitive ChR (λ_max_ = 460nm)^13^ in CA1 pyramidal neurons (Fig. 2E, see Methods for details). CheRiff-expressing pyramidal cells were readily spiking upon blue, but not orange-red illumination up to 10 mW/mm^2^ (Fig. 2E, S4). Conversely, red but not blue light triggered APs in somBiPOLES-expressing VIP neurons (Fig. 2E). Next, we recorded synaptic inputs from these two populations onto VIP-negative GABAergic neurons in stratum-oriens (Fig. 2F). As expected, blue light triggered EPSCs (CheRiff) and red light IPSCs (somBiPOLES), evident by their respective reversal potentials at 8.8 ± 10.4 mV and-71.4 ± 13.1 mV (Fig. 2F).

Next, we used BiPOLES to control cholinergic motor neurons in *C. elegans*. BiPOLES-linked mCerulean was visible at the nerve ring in the head part indicating correct expression and localization of the tool (Fig. 3A). Illumination with red light resulted in body-wall muscle contraction and effective body-shrinkage, consistent with motor neuron activation. Conversely, blue light triggered body extension, indicative of muscle relaxation and thus, cholinergic motor neuron inhibition (Fig. 3B). Maximal body length changes of +3% at 480 nm and-10% at 560-600 nm and reversal of the effect between 480-520 nm were consistent with the inhibitory and excitatory action spectrum of BiPOLES (Fig. 1E, 3B, S7) The light effects on body length required functional BiPOLES as light did not affect body length in the absence of all-*trans* retinal (ATR, Fig. 3B).

**Figure 3:**
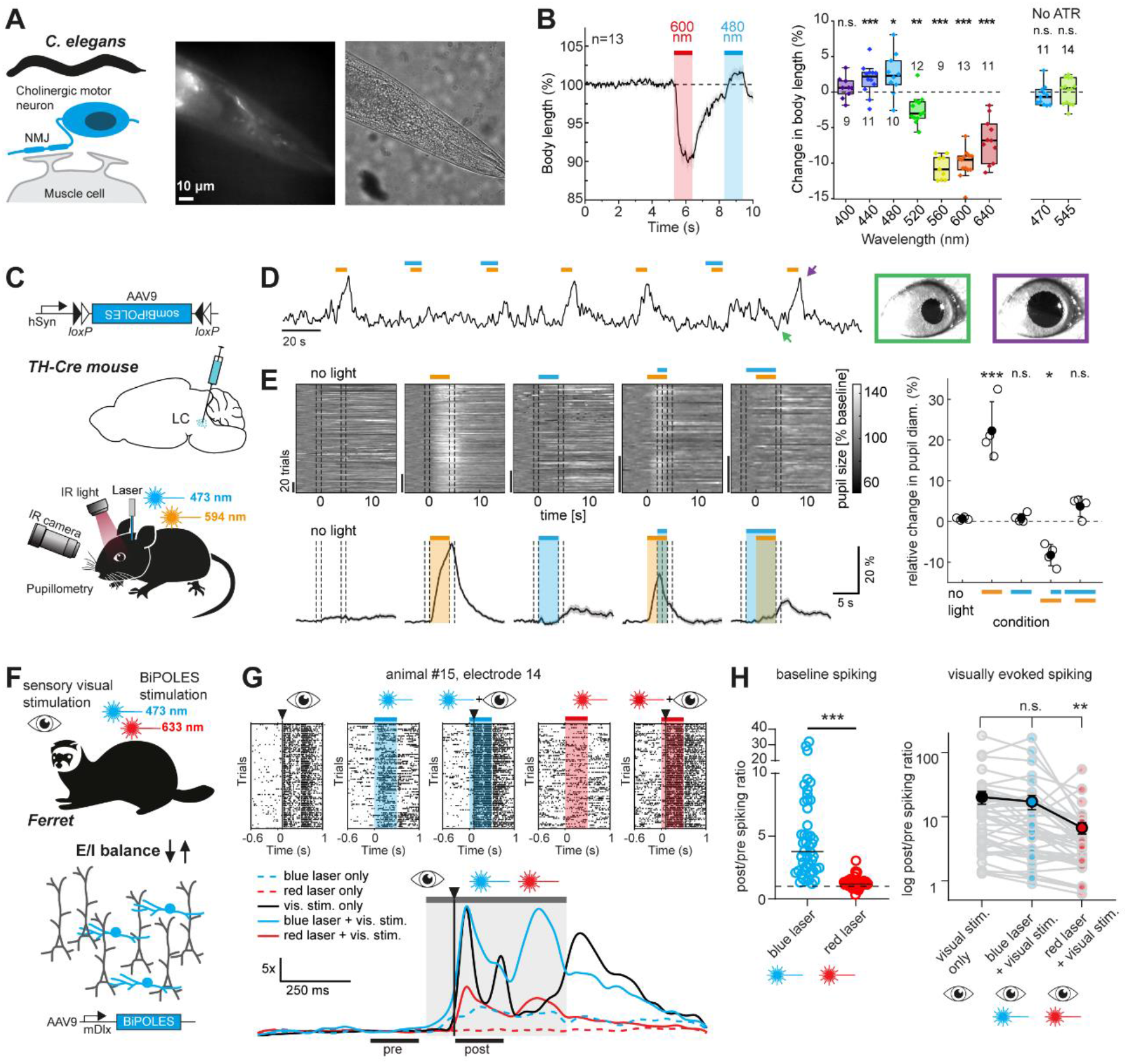
BiPOLES and somBiPOLES allow bidirectional modulation of neuronal activity *in vivo*. **(A)** BIPOLES expressed in cholinergic neurons of *C. elegans* enables bidirectional control of body contraction and relaxation. Left: Scheme of BiPOLES-expressing cholinergic motor neuron innervating a muscle cell. Right: Fluorescence and phase contrast micrographs showing expression of BiPOLES in the nerve ring. **(B)** Left: Temporal dynamics of relative changes in body length upon illumination with 600 and 480 nm light (1.1 mW/mm^2^, n = 13). Right: Spectral quantification of maximal change in body length (Box: median, 1^st^ – 3^rd^ quartile, whiskers: 1.5x inter quartile range, n = 9 - 14, *p < 0.05, **p < 0.01, ***p < 0.001). **(C)** Conditional expression of somBiPOLES in noradrenergic neurons of the mouse LC to modulate pupil dilation. **(D)** Relative pupil diameter in a single recording session. Orange and blue bars indicate time of illumination with 594 (orange) and 473 nm (blue), respectively. Arrows indicate positions of the two example images of the eye. **(E)** Quantification of normalized pupil size in one animal under various stimulation conditions for somBiPOLES as indicated. Top left: single trials. Bottom left: mean ± SEM. Dashed lines show regions used for quantification in the plot on the right. Right: quantification of relative pupil size (n = 4 mice, *p < 0.05, ***: p < 0.001). **(F)** Modulation of GABAergic neurons (blue) in ferret secondary visual cortex (area 18) with mDlx-BiPOLES. Red (633 nm) or blue (473 nm) laser light was used to (de-)activate interneurons with or without a preceding 10-ms visual flash (LED) to the ferret’s right eye. **(G)** Example neuronal spiking responses at one contact of the linear probe (∼900 µm depth) under indicated stimulation conditions Top: Raster-plots of the visual stimulus alone, blue laser (+visual), red laser (+visual) conditions. Bottom: Normalized to ‘pre’-phase averaged spike-density plot (sigma = 20 ms) of each indicated condition. Gray area: laser-on epoch; black vertical line: visual stimulus onset. Black horizontal lines indicate the 200 ms pre-and post-stim analysis epochs to compute the results in (H). Note the rate-increase after the onset of the blue laser before the onset of the visual stimulus and the reduced answer after red laser illumination. **(H)** Spike-rate ratio of pos-t vs pre-laser-stimulus epoch. Left: quantification of laser-mediated impact on baseline spiking rate (no visual stim.). Right: quantification of the spike-rate change of the same units during only visual and laser+visual stimulation. (n = 46 contacts showing visual responses from 3 animals, **p < 0.01, ***p < 0.001).

To extend the applications of BiPOLES, we generated various conditional and non-conditional viral vectors, in which the expression of the tandem protein is regulated by different promoters (see Methods, Figs. S8-9). Conditionally expressed in noradrenergic neurons of the Locus Coeruleus (LC) in mice, orange illumination of somBiPOLES reliably triggered pupil dilation, indicative of LC-mediated arousal^14^ (Fig. 3C-E, S10). Light-mediated pupil dilation was reverted immediately by additional blue light during the orange-light stimulation or suppressed altogether, when blue-light delivery started before orange-light application (Fig. 3D,E), suggesting that orange-light-induced spiking of noradrenergic neurons in LC was efficiently shunted. Thus, LC-neurons were bidirectionally controlled with somBiPOLES.

We hypothesized that targeting BiPOLES to GABAergic neurons enables bidirectional control of excitation/inhibition (E/I) balance. Thus, we generated a viral vector using the minimal *Dlx* promoter ^15^ (mDlx), verified mDlx-BiPOLES functionality *in vitro* (Fig. S8) and expressed it in GABAergic neurons in ferret secondary visual cortex to modulate E/I-balance during sensory processing (Fig. 3F).

Intracortical data obtained from linear probes and under isoflurane anesthesia provided evidence for modulation of cortical activity by shifts in E/I balance (Fig. 3G,H). Blue light led to an increase in baseline activity, consistent with deactivation of inhibitory neurons (Fig. 3G,H). Activation of GABAergic cells by red light did not further decrease the low cortical baseline activity, but significantly reduced cortical responses triggered by sensory stimuli (Fig. 3G,H). Although activating effects of blue light on evoked spiking were not significant in the average data, we obtained clear evidence in individual recordings that blue light could enhance late response components (Fig. 3G), confirming a disinhibitory effect. Overall, these data suggest that BiPOLES is efficient in bidirectional control of inhibitory mechanisms, demonstrating its applicability for the control of E/I shifts in the cortical microcircuit *in vivo*.

## Conclusion

In summary, BiPOLES is a tandem of a cation-and an anion-selective ChR that serves as a new optogenetic tool for balanced excitation and inhibition of the same neurons with red and blue light, respectively. In principle, multicistronic vectors encoding two opsins under a single promoter using an internal ribosomal entry site (IRES)^16^ or self-cleaving viral 2A peptide bridges allow expression of two rhodopsins at a fixed ratio from a single AAV vector^17,18^. However, these strategies do not ensure co-localized expression of the two rhodopsins since they may get differentially distributed in the plasma membrane and can have different degradation rates leading to variable stoichiometries of expression. In contrast, BiPOLES is a covalently linked fusion protein displaying a fixed expression of both opsins at a 1:1 stoichiometry anywhere in the membrane and membrane trafficking or degradation of both opsins occurs at an identical rate. Thus, excitatory and inhibitory currents are fixed to the same ratio in all expressing cells.

Compared to previous rhodopsin fusion constructs, that combined the excitatory blue-light-sensitive channelrhodopsin-2 (ChR2) with the orange-light-sensitive inhibitory ion pumps halorhodopsin (NphR) or bacteriorhosdopsin (bR)^10^, BiPOLES features a number of major advantages. First, combining two channels, rather than a pump and a channel, provides a more balanced quantum yield for the inhibitory and excitatory rhodopsin. Most importantly, both excitation and inhibition require only modest intensities of orange and blue light, respectively, whereas high light intensities are required for the ion pumps that only transport one charge per absorbed photon. Second, due to the use of two channels, BiPOLES-mediated photocurrents depend on the membrane polarity and thus, unlike with pumps, chloride or protons are not actively moved against their gradients, which can cause adverse side-effects^19^. Therefore, aside from dual-color inhibition and excitation, BiPOLES further allows to optically clamp the neuronal membrane voltage anywhere between the reversal potential of GtACR2 and Chrimson with a defined ratio of red/blue light. Third, compared to previous fusion constructs^10^, switching the color of the excitatory and inhibitory opsin restricts optical excitation in BiPOLES-expressing cells exclusively to the yellow/red spectrum. This modification enables scale-free and independent spiking of two neuronal populations in combination with a second, blue-light-sensitive ChR, expressed in the second population of neurons, as the blue-light-activated, inhibitory channel GtACR2 potently shunts Chrimson-mediated, blue-light-activated excitatory photocurrents. Fourth, compared to the first generation of tandem constructs, BiPOLES was optimized for membrane trafficking and especially the somBiPOLES variant shows strongly improved membrane trafficking in neurons, enabling reliable and potent optogenetic spiking and inhibition even in deep brain regions in vivo.

Since BiPOLES can be used to spike or inhibit the same population of neurons *in vivo*, a number of previously inaccessible questions can be addressed. For example, during extracellular recordings, BiPOLES may be useful for optogenetic identification (optotagging) with red light^20^ and optogenetic silencing of the same neurons. This allows verification of the identity of silenced neurons by their spiking profiles. Aside from bidirectional control of motor neurons, noradrenergic signaling in LC and GABAergic activity in neocortex, as demonstrated in this study, additional applications for BiPOLES could be bidirectional control of engram neurons^21^ to test both necessity and sufficiency of a particular set of neurons for memory retrieval or switching the valence of a particular experience by inhibiting or activating the same or even two distinct populations of neuromodulatory neurons. Due to its utility for a wide range of research questions and its applicability in numerous model systems, as demonstrated in this study, BiPOLES may fill a long-standing gap in the optogenetic toolbox.

## Methods

### Molecular Biology

For HEK-cell expression, the coding sequences of Chrimson (KF992060.1) and CsChrimson (KJ995863.2) from *Chlamydomonas noctigama*^1^, iC++ (Addgene #98165)^8^, Aurora (Addgene #98217)^7^, *Gt*ACR1 (KP171708) and *Gt*ACR2 (KP171709) from *Guillardia theta*^9^ were cloned together with mCerulean3^22^ and a trafficking signal (ts) from the Kir 2.1 channel^11^ into a pCDNA3.1 vector containing the original opsin tandem cassette^10^ with a linker composed of eYFP and the first 105 N-terminal amino acids of the rat gastric H+/K+-ATPase beta subunit (βHK, NM_012510.2), kindly provided by Sonja Kleinlogel (University of Bern, CH).

For neuronal expression, the insert consisting of GtACR2-ts-mCerulean3-βHK-Chrimson was cloned into an AAV2-backbone behind a human synapsin (hSyn) promoter (pAAV-hSyn-BiPOLES-mCerulean; Addgene #154944). A soma-targeted, membrane-trafficking optimized variant was generated by fusing an additional trafficking signal from the potassium channel Kv2.1^12^ to the C-terminus of Chrimson (pAAV-hSyn-somBiPOLES-mCerulean; Addgene #154945). For expression in GABAergic neurons, BiPOLES and somBiPOLES were cloned into an AAV2-backbone behind the minimal Dlx (mDlx) promoter^15^ resulting in pAAV-mDlx-BiPOLES-mCerulean (Addgene #154946) and pAAV-mDlx-somBiPOLES-mCerulean (Addgene #154947). For expression in projection neurons, somBiPOLES was cloned into an AAV2-backbone behind the minimal CaMKII promoter^23^ resulting pAAV-CaMKII-somBiPOLES-mCerulean (Addgene #154948). Double-floxed inverted open reading frame variants of BiPOLES and somBiPOLES were generated by cloning these inserts in antisense direction behind the Ef1alpha or hSyn promoter, flanked by two loxP and lox2272 sites (Ef1a-DIO-BiPOLES-mCerulean, Addgene #154949; hSyn-DIO-BiPOLES-mCerulean, Addgene #154950; hSyn-DIO-somBiPOLES-mCerulean, Addgene #154951). Note that in all constructs the mCerulean3-tag is fused between GtACR2-ts and βHK-Chrimson and therefore part of BiPOLES. We nonetheless chose to add “mCerulean” to the plasmid names to remind the reader of the presence of a cyan fluorophore in BiPOLES. BiPOLES stands for “<u>Bi</u>directional <u>P</u>air of <u>O</u>psins for <u>L</u>ight-induced <u>E</u>xcitation and <u>S</u>ilencing”. For reasons of political correctness, we chose this acronym over BREXIT (for: <u>B</u>lue-<u>R</u>ed <u>EXc</u>itation-Inhibition <u>T</u>ool).

### Patch-Clamp experiments in HEK293 cells

Patch-clamp experiments in HEK293 cells were prepared and performed as described previously^24^. Fusion constructs were expressed under the control of a CMV-promotor in HEK293 cells that were cultured in Dulbecco’s Modified Medium (DMEM) with stable glutamine (Biochrom, Berlin, Germany), supplemented with 10% (v/v) fetal bovine serum (FBS Superior; Biochrom, Berlin, Germany), 1 μM all-*trans* retinal, and 100 µg/ml penicillin/streptomycin (Biochrom, Berlin, Germany). Cells were seeded on poly-lysine coated glass coverslips at a concentration of 1 × 10^5^ cells/ml and transiently transfected using the FuGENE® HD Transfection Reagent (Promega, Madison, WI) two days before measurement.

Patch pipettes were prepared from borosilicate glass capillaries (G150F-3; Warner Instruments, Hamden, CT) using a P-1000 micropipette puller (Sutter Instruments, Novato, CA) and subsequently fire polished. Pipette resistance was between 1.2 and 2.5 MΩ. Single fluorescent cells were identified using an Axiovert 100 inverted microscope (Carl Zeiss, Jena, Germany). Monochromatic light (± 7 nm) was provided by a Polychrome V monochromator (TILL Photonics, Planegg, Germany). Light intensities were attenuated by a motorized neutral density filter wheel (Newport, Irvine, CA) for equal photon flux during action spectra recordings. Light pulses were controlled by a VS25 and VCM-D1 shutter system (Vincent Associates, Rochester, NY). Recordings were done with an AxoPatch 200B amplifier (Molecular Devices, Sunnyvale, CA), filtered at 2 kHz and digitized using a DigiData 1440A digitizer (Molecular Devices, Sunnyvale, CA) at a sampling rate of 10 kHz. The reference bath electrode was connected to the bath solution via a 140 mM NaCl agar bridge. Bath solutions contained 140 mM NaCl, 1 mM KCl, 1 mM CsCl, 2 mM CaCl_2_, 2 mM MgCl_2_ and 10 mM HEPES at pH_e_ 7.2 (with glucose added up to 310 mOsm). Pipette solution contained 110 mM NaGluconate, 1 mM KCl, 1 mM CsCl, 2 mM CaCl_2_, 2 mM MgCl_2_, 10 mM EGTA and 10 mM HEPES at pH_i_ 7.2 (glucose added up to 290 mOsm). All light intensities were measured in the object plane using a P9710 optometer (Gigahertz-Optik, Türkenfeld, Germany) and normalized to the water Plan-Apochromat 40×/1.0 differential interference contrast (DIC) objective illuminated field (0.066 mm^2^). The light intensity was 2.7 mW/mm^2^ at 650 nm, 3.5 mW/mm^2^ at 600 nm and 5.7 mW/mm^2^ at 490 nm. All electrical recordings were controlled by the pCLAMP™ software (Molecular Devices, Sunnyvale, CA). All whole-cell recordings had a membrane resistance of at least 500 MΩ (usual >1 GΩ) and an access resistance below 10 MΩ.

### Organotypic hippocampal slice culture preparation and transgene delivery

Organotypic hippocampal slices were prepared from Wistar rats or VIP-IRES-Cre mice (Jackson-No. 031628) at post-natal day 5-7 as described^25^. Briefly, dissected hippocampi were cut into 350 μm slices with a tissue chopper and placed on a porous membrane (Millicell CM, Millipore). Cultures were maintained at 37°C, 5% CO_2_ in a medium containing 80% MEM (Sigma M7278), 20% heat-inactivated horse serum (Sigma H1138) supplemented with 1 mM L-glutamine, 0.00125% ascorbic acid, 0.01 mg/ml insulin, 1.44 mM CaCl_2_, 2 mM MgSO_4_ and 13 mM D-glucose. No antibiotics were added to the culture medium.

For transgene delivery in organotypic slices, individual CA1 pyramidal cells were transfected by single-cell electroporation between DIV 14-16 as previously described^26^. The plasmids pAAV-hSyn-BiPOLES-mCerulean, pAAV-hSyn-somBiPOLES-mCerulean, and pAAV-hSyn-Chrimson-mCerulean were diluted to 5 ng/µl in K-gluconate-based solution consisting of (in mM): 135 K-gluconate, 10 HEPES, 4 Na_2_-ATP, 0.4 Na-GTP, 4 MgCl_2_, 3 ascorbate, 10 Na_2_-phosphocreatine (pH 7.2). A plasmid encoding hSyn-mKate2 (50 ng/µl) was co-electroporated and served as a morphology marker. An Axoporator 800A (Molecular Devices) was used to deliver 50 hyperpolarizing pulses (−12 V, 0.5 ms) at 50 Hz. During electroporation slices were maintained in pre-warmed (37°C) HEPES-buffered solution (in mM): 145 NaCl, 10 HEPES, 25 D-glucose, 2.5 KCl, 1 MgCl_2_ and 2 CaCl_2_ (pH 7.4, sterile filtered). In some cases, slice cultures were transduced with adeno-associated virus (see Table 1 for details) at DIV 3-5 as described^27^. Briefly, the different rAAVs were locally injected into the CA1 region using a Picospritzer (Parker, Hannafin) by a pressurized air pulse (2 bar, 100 ms) expelling the viral suspension into the slice. During virus infection, membranes carrying the slices were kept on pre-warmed HEPES-buffered solution.

**Table 1.** List of adeno-associated viral vectors used for experiments in organotypic hippocampal slices. Virus were transduced at the indicated titers.

| Adeno-associated virus (AAV9) | Titer used for transduction of hippocampal organotypic slice cultures (vg/ml) | Addgene plasmid reference |
| --- | --- | --- |
| hSyn-DIO-somBiPOLES-mCerulean | $3.4 \times 10^{13}$ | 154951 |
| mDlx-BiPOLES-mCerulean | $2.8 \times 10^{13}$ | 154946 |
| CaMKIIa(0.4)-somBiPOLES-mCerulean | $2.5 \times 10^{13}$ | 154948 |
| CaMKIIa(0.4)-DO-CheRiff-ts-mScarlet-ER | $8.15 \times 10^{11}$ | n.a. |
| mDlx-H2B-EGFP | $2.8 \times 10^{10}$ | n.a. |

### Slice culture two-photon microscopy

Neurons in organotypic slice cultures (DIV 19-21) were imaged with two-photon microscopy to characterize their morphology and the subcellular localization of BiPOLES and somBiPOLES. The custom-built two-photon imaging setup was based on an Olympus BX-51WI upright microscope upgraded with a multiphoton imaging package (DF-Scope, Sutter Instrument), and controlled by ScanImage 2017b software (Vidrio Technologies). Fluorescence was detected through the objective (Olympus LUMPLFLN 60XW, 1.0 NA, or Leica HC FLUOTAR L 25x/0.95 W VISIR) and through the oil immersion condenser (numerical aperture 1.4, Olympus) by two pairs of GaAsP photomultiplier tubes (Hamamatsu, H11706-40). Dichroic mirrors (560 DXCR, Chroma Technology) and emission filters (ET525/70m-2P, ET605/70m-2P, Chroma Technology) were used to separate cyan and red fluorescence. Excitation light was blocked by short-pass filters (ET700SP-2P, Chroma Technology). A tunable Ti:Sapphire laser (Chameleon Vision-S, Coherent) was set to 810 nm to excite mCerulean on BiPOLES and somBiPOLES. An Ytterbium-doped 1070-nm pulsed fiber laser (Fidelity-2, Coherent) was used at 1070 nm to excite the volume filler mKate2. Maximal intensity projections of z-stacks were generated with Fiji ^28^

### Slice culture electrophysiology

At DIV 19-21, whole-cell patch-clamp recordings of transfected or virus-infected CA1 pyramidal or GABAergic neurons were performed. Experiments were done at room temperature (21-23°C) under visual guidance using a BX 51WI microscope (Olympus) equipped with Dodt-gradient contrast and a Double IPA integrated patch amplifier controlled with SutterPatch software (Sutter Instrument, Novato, CA). Patch pipettes with a tip resistance of 3-4 MΩ were filled with intracellular solution consisting of (in mM): 135 K-gluconate, 4 MgCl_2_, 4 Na_2_-ATP, 0.4 Na-GTP, 10 Na_2_-phosphocreatine, 3 ascorbate, 0.2 EGTA, and 10 HEPES (pH 7.2). Artificial cerebrospinal fluid (ACSF) consisted of (in mM): 135 NaCl, 2.5 KCl, 2 CaCl_2_, 1 MgCl_2_, 10 Na-HEPES, 12.5 D-glucose, 1.25 NaH_2_PO_4_ (pH 7.4). In experiments where synaptic transmission was blocked, 10 µM CPPene, 10 µM NBQX, and 100 µM picrotoxin (Tocris, Bristol, UK) were added to the recording solution. In experiments analyzing synaptic inputs onto O-LM interneurons, ACSF containing 4 mM CaCl_2_ and 4 mM MgCl_2_ was used to reduce the overall excitability. Measurements were corrected for a liquid junction potential of-14,5 mV. Access resistance of the recorded neurons was continuously monitored and recordings above 30 MΩ were discarded. A 16 channel LED light engine (CoolLED pE-4000, Andover, UK) was used for epifluorescence excitation and delivery of light pulses for optogenetic stimulation (ranging from 385 to 635 nm). Light intensities were measured in the object plane with a 1918 R power meter equipped with a calibrated 818 ST2 UV/D detector (Newport, Irvine CA) and divided by the illuminated field of the Olympus LUMPLFLN 60XW objective (0.134 mm^2^).

In current-clamp experiments holding current was injected to maintain CA1 cells near their resting membrane potential (−75 to-80 mV). To compare the spiking probability of somBiPOLES and Chrimson under illumination with light of different wavelengths (470, 595 and 635 nm), a train of 20 light pulses (5 ms pulse duration) was delivered at 5 Hz. For each wavelength, light intensities from 0.1 to 100 mW/mm^2^ were used.

To measure the ability of BiPOLES and somBiPOLES to shift the rheobase upon blue-light illumination, depolarizing current ramps (from 0–100 to 0–900 pA) were injected into CA1 neurons in the dark and during illumination with 490 nm light at intensities ranging from 0.001 to 10 mW/mm^2^. The injected current at the time of the first spike was defined as the rheobase. The relative change in the number of ramp-evoked action potentials (APs) was calculated counting the total number of APs elicited during the 9 current ramp injections (from 0–100 to 0–900 pA) for each light intensity and normalized to the number of APs elicited in the absence of light. Statistical significance was calculated using Friedman test.

To assess the capability of BiPOLES and somBiPOLES as dual-color neuronal excitation and silencing tools, alternating pulses of red (635 nm, 20 ms, 10 mW/mm^2^), blue (490 nm, 100 ms, 10 mW/mm^2^) and a combination of these two (onset of blue light 40 ms before red light) were delivered to elicit and block action potentials.

For independent optogenetic activation of two distinct populations of neurons, organotypic slice cultures from VIP-Cre mice were transduced with 2 adeno-associated viral vectors: 1, a double-floxed inverted open reading frame (DIO) construct encoding somBiPOLES (hSyn-DIO-somBiPOLES-mCerulean, see Table 1 for details) to target VIP-positive interneurons, and 2, a double-floxed open reading frame (DO) construct encoding CheRiff (hSyn-DO-CheRiff-ts-mScarlet-ER, see Table 1 for details) to target CA1 pyramidal neurons and exclude expression in VIP-positive cells. Synaptic input from these two populations was recorded in VIP-negative stratum-oriens GABAergic neurons (putative O-LM cells). In CA1, O-LM neurons receive innervation both from local CA1 pyramidal cells and VIP-positive GABAergic neurons ^29^. To facilitate identification of putative GABAergic post-synaptic neurons in stratum oriens, slices were transduced with an additional rAAV encoding mDlx-H2B-EGFP. In the absence of synaptic blockers light-evoked EPSCs and IPSCs were recorded while holding the postsynaptic cell at different membrane potentials (from-80,-65,-55,-45 and 6 mV) in whole-cell voltage clamp mode. A blue (460 nm, 0.03-84.0 mW/mm^2^) and a red (635 nm, 6.0 – 97.0 mW/mm^2^) light pulse were delivered 500 ms apart from each other through a Leica HC FLUOTAR L 25x/0.95 W VISIR objective.

In experiments determining the light intensity threshold needed to spike CA1 cells with BiPOLES, somBiPOLES, Chrimson and CheRiff across different wavelengths, 470, 525, 595 and 635 nm light ramps going from 0 to 10 mW/mm^2^ over 1 s were delivered in current-clamp mode. In the case of BiPOLES and somBiPOLES the blue light ramp went up to 100 mW/mm^2^ to rule out that very high blue light intensities might still spike neurons. The light intensity value at the time of the first spike was defined as the light intensity threshold (in mW/mm^2^) needed to evoke action potential firing.

For photocurrent density measurements in voltage-clamp mode CA1 cells were held at −75 or −55 mV to detect inward (cationic) or outward (anionic) currents elicited by red (635 nm, 20 ms, 1 and 10 mW/mm^2^) and blue light (490 nm, 100 ms, 10 mW/mm^2^), respectively. For each cell, the peak photocurrent amplitude (in pA) was divided by the cell membrane capacitance (in pF) which was automatically recorded by the Sutterpatch software in voltage-clamp mode (V_hold_ =−75 mV).

To optically clamp the neuronal membrane potential using somBiPOLES, simultaneous illumination with blue and orange light at varying ratios was used. In current-clamp experiments, 470 and 595 nm light ramps (5 s) of opposite gradient (1 to 0 mW/mm^2^ and 0 to 1 mW/mm^2^, respectively) were applied. To reveal the slow change in membrane voltage during the ramps protocol, spikes elicited at high 595/470 nm ratios were removed by median-filtering the voltage traces.

### Transgenic *C. elegans* lines and transgenes

For expression in cholinergic neurons of *C. elegans*, BiPOLES (GtACR2::ts::mCerulean3::βHK::Chrimson) was subcloned into the p*unc*-17 vector RM#348p (a gift from Jim Rand) via Gibson Assembly based on the plasmid CMV_GtACR2_mCerulean_βHK_Chrimson, using the restriction enzyme *Nhe*I and the primers ACR2_Chrimson_fwd (5’-attttcaggaggacccttggATGGCATCACAGGTCGTC-3’) and ACR2_Chrimson_rev (5’-ataccatggtaccgtcgacgTCACACTGTGTCCTCGTC-3’), resulting in the construct pAB26. The respective transgenic strain ZX2586 (wild type; *zxEx1228[punc-17::GtACR2::ts::mCerulean3::βHK::Chrimson;* p*elt-2::GFP]*), was generated via microinjection ^30^ of both 30 ng/µl plasmid and co-marker plasmid DNA p*elt-2*::GFP. Animals were cultivated on nematode growth medium (NGM), seeded with *E. coli* OP-50 strain, in 6 cm petri dishes. To obtain functional rhodopsins in optogenetic experiments, the OP-50 bacteria were supplemented with all-*trans* retinal ATR (0.25 μl of a 100 mM stock (in ethanol) mixed with 250 μl OP-50 bacterial suspension).

### *C. elegans* stimulation and behavioral experiments

For body-length measurements, L4 stage transgenic animals were cultivated on ATR plates overnight. Video analysis of light-stimulation protocols provided information on depolarized and hyperpolarized states, based on contracted or relaxed body-wall muscles (BWMs)^31^. Prior to experiments, animals were singled on plain NGM plates to avoid imaging artefacts. They were manually tracked with an Axio Scope.A1 microscope (Zeiss, Germany), using a 10x objective (Zeiss A-Plan 10x/0,25 Ph1 M27) and a Powershot G9 digital camera (Canon, USA). For light-stimulation of BiPOLES, transgenic worms were illuminated with 1 or 5 s light pulses at 1.1 mW/mm^2^ of different wavelengths as indicated in Fig. 3B, S7 (monochromatic light source, Polychrome V, Till Photonics), controlled via an Arduino-driven shutter (Sutter Instrument, USA). Videos were processed and analyzed using a custom written MATLAB script^32^ (MathWorks, USA). For the analysis of data, the animals’ body length was normalized to the recording period prior to illumination.

### Modulation of arousal via control of noradrenergic neurons in the mouse locus coeruleus (LC)

#### Animals

All procedures were in agreement with the German national animal care guidelines and approved by the Hamburg state authority for animal welfare (BGV; license 33/19) and the animal welfare officer of the University Medical Center Hamburg-Eppendorf. Experiments were performed on mice of either sex between 2.5 and 4 months of age at the start of the experiment. Mice were obtained from The Jackson Laboratory, bred and maintained at our own colony (12/12h light-dark cycle, 22°C room temperature, ∼40% relative humidity, food and water ad libitum). Transgenic mice expressing Cre recombinase in tyrosine hydroxylase positive neurons (TH-Cre, Stock No: 008601) ^33^ were injected with a suspension of rAAV9 viral particles encoding hSyn-DIO-somBiPOLES (see table 1) to target noradrenergic neurons in the locus coeruleus. Control experiments were performed in non-injected wildtype littermates.

#### Virus injection and implantation of optic fibers

General anesthesia and analgesia was achieved by intraperitoneal injections of midazolam/medetomidine/fentanyl (5.0/0.5/0.05 mg/kg, diluted in NaCl). After confirming anesthesia and analgesia by absence of the hind limb withdrawal reflex, the scalp of the animal was trimmed and disinfected with Iodide solution (Betaisodona; Mundipharma, Germany). The animal was placed on a heating pad to maintain body temperature, fixed in a stereotactic frame, and the eye ointment (Vidisic; Bausch + Lomb, Germany) was applied to prevent drying of the eyes. To bilaterally access the LC, an incision (1-2 cm) was made along the midline of the scalp, the skull was cleaned, and small craniotomies were drilled-5.4 mm posterior and ± 1 mm lateral to Bregma. 0.4 µl of virus suspension were injected into each LC (−3.6 mm relative to Bregma) at a speed of ∼100-200 nl/min using a custom-made air pressure system connected to a glass micropipette. After each injection, the micropipette was left in place for a minimum of 5 minutes before removal. After virus injection, cannulas housing two ferrule-coupled optical fibers (200 µm core diameter, 0.37 NA, 4 mm length) spaced 2 mm apart (TFC_200/245-0.37_4mm_TS2.0_FLT; Doric Lenses, Canada) were inserted just above the injection site to a depth of-3.5 mm relative to Bregma using a stereotactic micromanipulator. The implant, as well as a headpost for animal fixation during the experiment, were fixed to the roughened skull using cyanoacrylate glue (Pattex; Henkel, Germany) and dental cement (Super Bond C&B; Sun Medical, Japan). The incised skin was glued to the cement to close the wound. Anesthesia was antagonized by intraperitoneally injecting a cocktail of atipamezole/flumazenil/buprenorphine (2.5/0.5/0.1 mg/kg, diluted in NaCl). Carprofen was given subcutaneously for additional analgesia and to avoid inflammation. In addition, animals received meloxicam mixed into softened food for 3 days after surgery.

#### Optogenetic stimulation

4-6 weeks after surgery, mice were habituated to head fixation and placement in a movement-restraining plastic tube for at least one session. Bilateral optogenetic stimulation of LC neurons was achieved by connecting the fiber implant to a 1×2 step-index multimode fiber optic coupler (200 µm core diameter, 0.39 NA; TT200SL1A, Thorlabs, Germany) in turn connected to a laser combiner system (LightHUB; Omicron, Germany) housing a 473 nm (LuxX 473-100; Omicron, Germany) and a 594 nm diode laser (Obis 594 nm LS 100 mW; Coherent, Germany) for activation the GtACR2 and Chrimson components of somBiPOLES, respectively. Coupling to the implant was achieved with zirconia mating sleeves (SLEEVE_ZR_1.25; Doric lenses, Canada) wrapped with black tape to avoid light emission from the coupling interface. Following a habituation period of ∼3 min after placing in the setup, stimuli were generated and presented using custom-written MATLAB scripts (MathWorks, US) controlling a NI-DAQ-card (PCIe-6323; National Instruments, US) to trigger the lasers via digital input channels. For activation of Chrimson, pulse trains (594 nm, ∼10 mW at each fiber end, 20 ms pulse duration, 20 Hz repetition rate) of 4 s duration were presented, while GtACR2 was activated by constant illumination (473 nm, ∼10 mW at each fiber end) of 2-6 seconds duration. 30-40 trials of 473 nm pulses, 594 nm pulse trains, and combinations thereof, were presented at an inter-train-interval of 20-30 seconds in each session.

#### Data acquisition

A monochrome camera (DMK 33UX249; The Imaging Source, Germany) equipped with a macro objective (TMN 1.0/50; The Imaging Source, Germany) and a 780 nm long-pass filter (FGL780; Thorlabs, Germany) was pointed towards one eye of the mouse. Background illumination was provided with an infrared spotlight (850 nm), while a UV LED (395 nm; Nichia, Japan) was adjusted to maintain pupil dilation of the mouse at a moderate baseline level. Single frames were triggered at 30 Hz by an additional channel of the NI-DAQ-card that controlled optogenetic stimulation, and synchronization was achieved by simultaneous recording of all control voltages and their corresponding timestamps.

#### Data analysis

Pupil diameter was estimated using a custom-modified, MATLAB-based algorithm developed by McGinley et al ^34^. In short, an intensity threshold was chosen for each recording to roughly separate between pupil (dark) and non-pupil (bright) pixels. For each frame, a circle around the center of mass of putative pupil pixels and with an area equivalent to the amount of pupil pixels was then calculated, and putative edge pixels were identified by canny edge detection. Putative edge pixels that were more than 3 pixels away from pixels below the threshold (putative pupil) or outside an area of ± 0.25-1.5 times the diameter of the fitted circle were neglected. Using least-squares regression, an ellipse was then fit on the remaining edge pixels, and the diameter of a circle of equivalent area to this ellipse was taken as the pupil diameter. Noisy frames (e.g. no visible pupil due to blinking or blurry pupil images due to saccades of the animal) were linearly interpolated, and the data was low-passed filtered (< 3 Hz; 3^rd^ order Butterworth filter). Pupil data was segmented from 5 s before to 15 s after onset of each stimulus and normalized to the median pupil diameter of the 5 s preceding the stimulus onset, before individual trials were averaged. Randomly chosen segments of pupil data of the same duration served as a control. The difference in median pupil diameter one second before and after stimulation (as indicated in Fig. 3E) was used to calculate potential changes in pupil diameter for each condition. Statistical significance was calculated using one-way analysis of variance and post-hoc multiple comparison tests.

### In-vivo recordings from ferret visual cortex

Data were collected from 3 adult female ferrets (*Mustela putorius*). Details on the surgical procedures have been reported earlier ^35,36^. All experiments were approved by the independent Hamburg state authority for animal welfare (BUG Hamburg) and were performed in accordance with the guidelines of the German Animal Protection Law.

For injection of rAAV9 viral particles encoding mDlx-BiPOLES-mCerulean (see table 1) animals were anesthetized with an injection of ketamine (15 mg/kg), medetomidine (0.02 mg/kg), midazolam (0.5 mg/kg) and atropine (0.15 mg/kg). Subsequently, they were intubated and respirated with a mixture of 70:30 N_2_/O_2_ and 1-1.5% isoflurane. A cannula was inserted into the femoral vein to deliver a bolus injection of enrofloxacin (15 mg/kg) and rimadyl (4 mg/kg) and, subsequently, continuous infusion of 0.9% NaCl and fentanyl (0.01 mg/kg/h). Body temperature, heart rate and end-tidal CO_2_ were constantly monitored throughout the surgery. The temporalis muscle was folded back, such that a small craniotomy (ø: 2.5mm) could be performed over the left posterior cortex and the viral construct was slowly (1µl/10min) injected into secondary visual cortex (area 18). The excised piece of bone was put back in place and fixed with tissue-safe silicone (Kwikcast; WPI). Also, the temporalis muscle was returned to its physiological position and the skin was closed. After the surgery the animals received preventive analgesics (Metacam, 0.1 mg) and antibiotics (Enrofloxacin, 15 mg/kg) for ten days.

After an expression period of at least 4 weeks, recordings of cortical signals were carried out under isoflurane anesthesia. Anesthesia induction and maintenance were similar to the procedures described above, except for a tracheotomy performed to allow for artificial ventilation of the animal over an extended period. The i.v. infusion was supplemented with pancuronium bromide (6 µg/kg/h) to prevent slow ocular drifts. Before fixing the animal’s head in the stereotaxic frame, a local anesthetic (Lidocaine, 10%) was applied to the external auditory canal. To keep the animal’s head in a stable position throughout the placement of recording electrodes and the measurements, a headpost was fixed with screws and dental acrylic to the frontal bone of the head. Again, the temporalis muscle was folded back and a portion of the cranial bone was resected. The dura was removed before introducing an optrode with 32 linearly distributed electrodes (A1×32-15mm-50(100)-177, NeuroNexus Technologies) into the former virus-injection site (area 18). The optrode was manually advanced via a micromanipulator (David Kopf Instruments) under visual inspection until the optic fiber was positioned above the pial surface and the uppermost electrode caught a physiological signal, indicating that it had just entered the cortex.

During electrophysiological recordings the isoflurane level was maintained at 0.7%. To ensure controlled conditions for sensory stimulation, all experiments were carried out in a dark, sound-attenuated anechoic chamber (Acoustair, Moerkapelle, Netherlands). Visual stimuli were created via an LED placed in front of the animal’s eye. In separate blocks, 150 laser stimuli of different colors (‘red’, 633nm LuxXplus and ‘blue’, 473nm LuxXplus, LightHub-4, Omicron) were applied through the optrode for 500 ms, each, at a variable interval of 2.5-3 seconds. Randomly, 75 laser stimuli were accompanied by a 10ms LED-flash, starting 100ms after the respective laser onset. For control, one block of 75 LED-flashes alone were presented at comparable interstimulus intervals.

Electrophysiological signals were sampled with an AlphaLab SnR recording system (Alpha Omega Engineering, Nazareth, Israel) or with a self-developed neural recording system based on INTAN digital head-stages (RHD2132, Intantech). Signals recorded from the intracortical laminar probe were band-pass filtered between 0.5 Hz and 7.5 kHz and digitized at 22-44 kHz or 25 kHz, respectively. All analyses of neural data presented in this study were performed offline after the completion of experiments using custom-written MATLAB scripts (MathWorks). To extract multiunit spiking activity (MUA) from broadband extracellular recordings, we high-pass filtered signals at 500Hz and detected spikes at negative threshold (>3.5 SD) ^37^.

### Data availability

The datasets generated during and/or analyzed during the current study are available from the corresponding author on reasonable request. The computer code generated to acquire or analyze data during the current study is available from the corresponding author on reasonable request, as well.

## Acknowledgements

We thank Stefan Schillemeit, Sandra Augustin and Tharsana Tharmalingam for excellent technical assistance, Mathew McGinley and Peter Murphy for help with pupil analysis, Sonja Kleinlogel for providing plasmids carrying the original opsin tandem cassette and Jonas Wietek for providing ACR plasmids and highly appreciated discussions at an early phase of the project. Ingke Braren of the UKE Vector Facility produced AAV vectors. This work was supported by the German Research Foundation, DFG (SPP1926, FOR2419/P6, SFB963/B8 to J.S.W., SFB936/A2 and SPP2041/EN533/15-1 to A.K.E., SPP1926 to P.H., SFB807/P11 to A.B. & A.G.) and the European Research Council (ERC2016-StG-714762 to J.S.W., Stardust to P.H.). Peter Hegemann is a Hertie Professor and supported by the Hertie Foundation.

## Author contributions

Conceptualization, J.V., S.R.R., P.H., J.S.W.; Methodology J.V., S.R.R., F.P., A.D., J.A., A.K.E., A.G., P.H., J.S.W.; Experimentation, J.V., S.R.R., F.P., A.D., A.B. N.Z., J.A.; Analysis, J.V., S.R.R., F.P., A.D., A.B. N.Z., J.A.; Writing, J.V., S.R.R., A.K.E., A.G., J.S.W.; Supervision, A.K.E., A.G., P.H., J.S.W.; Funding Acquisition, A.B., A.K.E., A.G., P.H., J.S.W.

## Notes

### Competing Interest Statement

The authors have declared no competing interest.

### Summary of Updates

Author affiliations were wrongly indicated in the original PDF-version of the preprint.

