## Supplemental figures for "BiPOLES: a tool for bidirectional dual-color optogenetic control of neurons"

### Supplementary Figures

**Fig. S1**

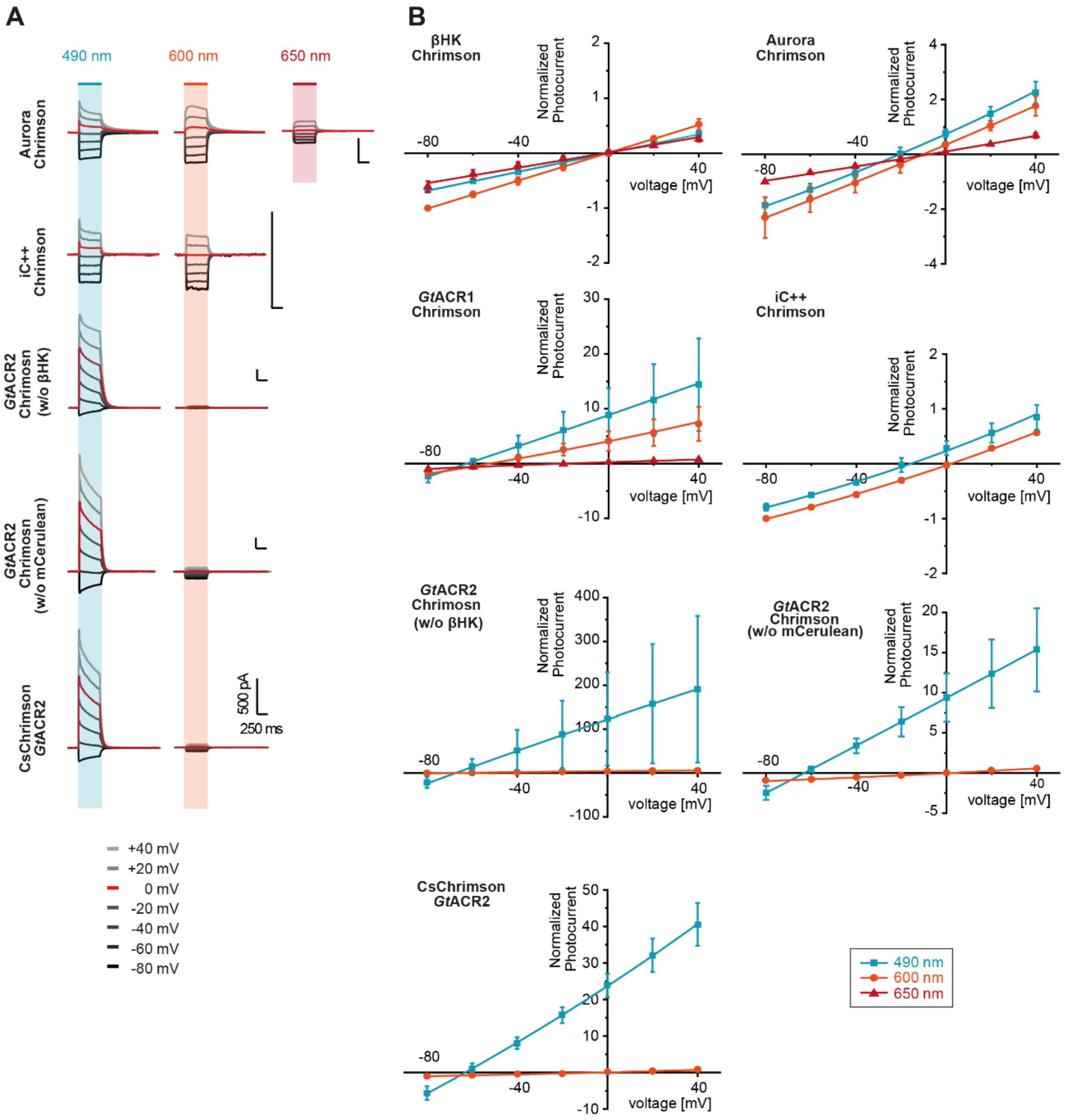


**Figure S1: Biophysical characterization of various ACR-Chrimson tandem variants. (A)** Representative photocurrent traces of Aurora-ts-eYFP-βHK-Chrimson, iC++-ts-eYFP-βHK-Chrimson, *Gt*ACR2-ts-mCerulean-Chrimson, *Gt*ACR2-ts-βHK-Chrimson and CsChrimson-ts-mCerulean-βHK-*Gt*ACR2 from whole-cell patch clamp recordings of HEK293 cells evoked by 490 nm, 600 nm and 650 nm illumination. Extra- and intracellular ionic conditions are indicated in Fig. 1A. **(B)** Normalized peak photocurrents of the fusion constructs shown in (A) as well as βHK-Chrimson-mCerulean and *Gt*ACR1-ts-mCerulean-βHK-Chrimson at different membrane voltages during illumination with either 490 nm, 600 nm or 650 nm light (Mean ± SD; n = 2 - 8; normalized to the peak photocurrent at -80 mV and 600 nm illumination (for βHK-Chrimson, iC++-Chrimson, *Gt*ACR2-Chrimson (w/o mCer), CsChrimson-*Gt*ACR2) or 650 nm illumination (for Aurora-Chrimson, *Gt*ACR1-Chrimson).

Fig. S2

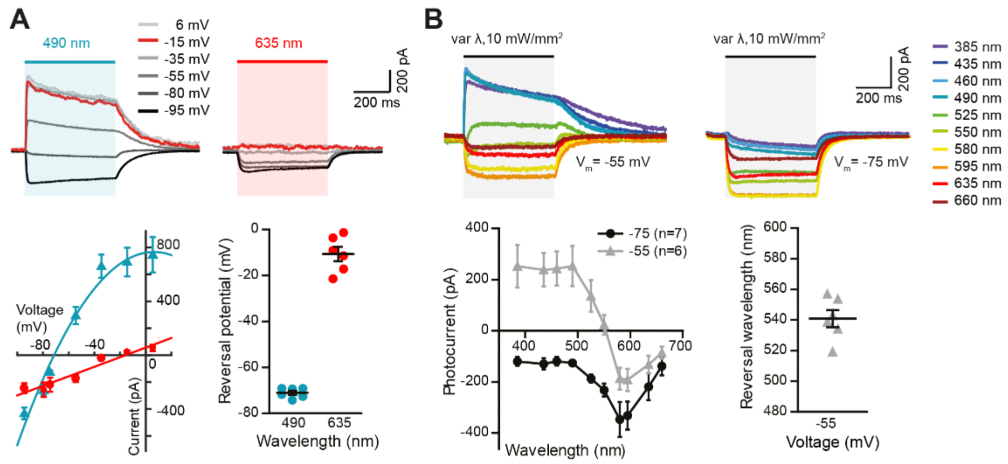


**Figure S2: Biophysical characterization of BiPOLES in CA1 pyramidal neurons. (A)** Top: Representative photocurrent traces of BiPOLES in CA1 pyramidal neurons at indicated membrane voltages upon illumination with 490 nm or 635 nm. Bottom: Quantification of photocurrent-voltage relationship (left) and photocurrent reversal potential (right) under 490 nm and 635 nm illumination (mean ± SEM, n = 6). Reversal of 490 and 635 nm photocurrents close to chloride and proton Nernst potentials indicates functional expression of BiPOLES in CA1 pyramidal neurons. **(B)** Top: Representative photocurrent traces of BiPOLES in CA1 pyramidal neurons upon illumination with different wavelengths and equal photon flux at membrane voltages above (left) and below (right) the chloride Nernst potential. Bottom: Quantification of photocurrents (left) and the photocurrent reversal wavelength at -55 mV (mean ± SEM, n = 6 - 7). Similar to HEK-cell measurements, inward and outward photocurrents were evoked with 635 nm and 490 nm at a membrane voltage between the chloride and proton Nernst potentials, respectively, indicative of independently evoked Chrimson- and *Gt*ACR2-photocurrents.

**Fig. S3**

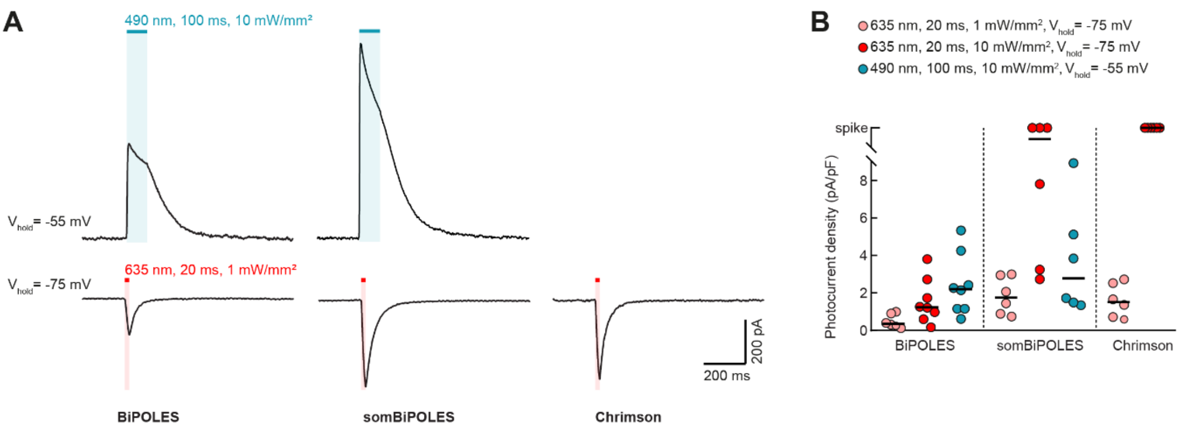


**Figure S3: Photocurrent density analysis in CA1 pyramidal cells. (A)** Representative photocurrent traces measured in BiPOLES-, somBiPOLES- or Chrimson-expressing CA1 pyramidal neurons. For all three constructs, photocurrents evoked by a 635 nm light pulse (20 ms, 1 mW/mm$^2$) were recorded at a membrane voltage of -75 mV. For BiPOLES and somBiPOLES, the *Gt*ACR2-photocurrent evoked by a 490 nm light pulse (100 ms, 10 mW/mm$^2$) was recorded in addition at a membrane voltage of -55 mV. **(B)** Quantification of photocurrent densities evoked under the indicated conditions. Note that photocurrent densities were strongly enhanced for somBiPOLES compared to BiPOLES and similar to Chrimson (black horizontal lines: medians, n = 6 - 8).

**Fig. S4**

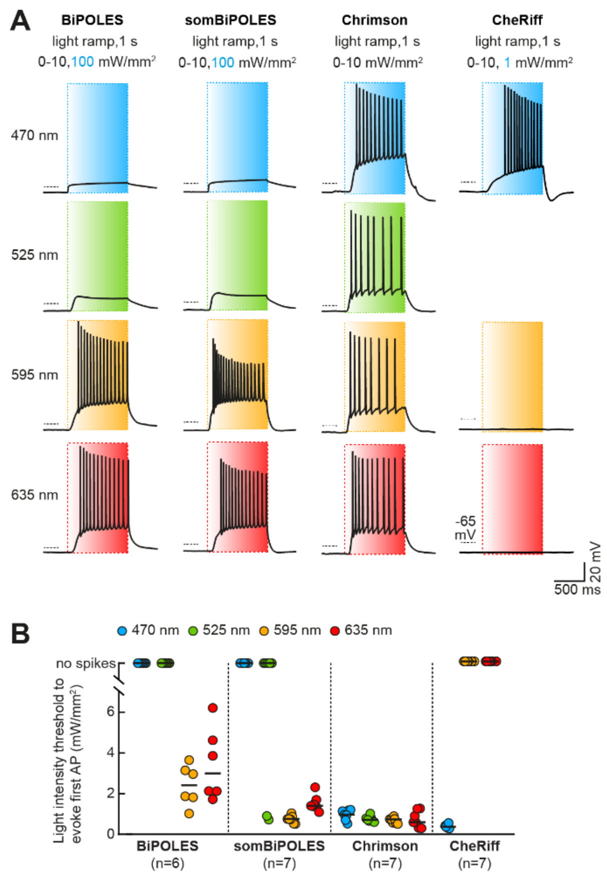


**Figure S4: Spectral quantification of action potential threshold for BiPOLES, somBiPOLES, Chrimson and CheRiff. (A)** Representative membrane voltage traces measured in BiPOLES-, somBiPOLES-, Chrimson-, or CheRiff-expressing CA1 pyramidal neurons. In IC experiments, light ramps were applied as indicated. The intensity was ramped linearly from 0 to 10 mW/mm$^2$ over 1 s, except for BiPOLES and somBiPOLES-expressing cells, where 470-nm ramps were ranging to 100 mW/mm$^2$ to rule out the possibility that high-intensity blue light might still evoke action potentials. For CheRiff, which has the absorption peak at 470 nm, blue-light ramps ranging to 1 mW/mm$^2$ were sufficient. **(B)** Quantification of the light intensity threshold at which the first action potential was evoked. 470-nm light up to 100 mW/mm$^2$ did not evoke action potentials in BiPOLES or somBiPOLES-expressing cells. somBiPOLES-expressing cells showed a reduced action potential threshold compared to BiPOLES for 595 and 635 nm indicating higher light sensitivity. The light-intensity threshold for action potential firing with 595 nm was similar between somBiPOLES- and Chrimson-expressing neurons, suggesting that with somBiPOLES action potentials can be evoked with high efficacy by orange-red-light, while action potentials are not triggered by blue light of any intensity (black horizontal lines: medians, n = 6 - 7).

**Fig. S5**

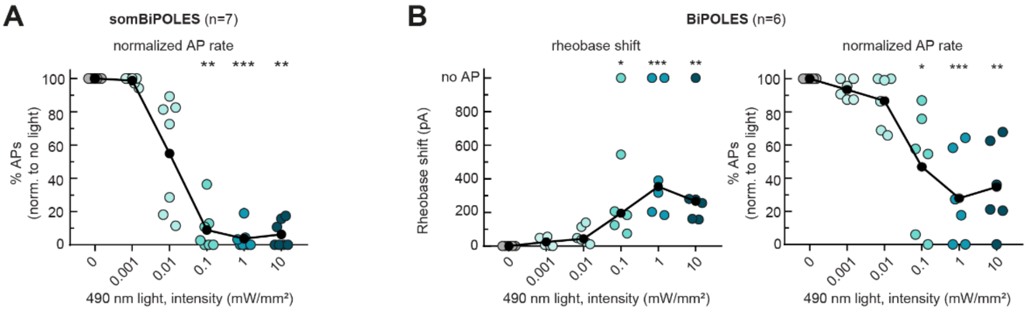


**Figure S5: Quantification of BiPOLES- and somBiPOLES-mediated neuronal silencing.** IC characterization of BiPOLES- and somBiPOLES mediated neuronal silencing. Current ramps (from 0–100 to 0–900 pA) were injected into BiPOLES- or somBiPOLES expressing CA1 pyramidal cell to induce action potentials (APs). **(A)** Relative change in the number of ramp-evoked action potentials in somBiPOLES-expressing neurons upon illumination with blue light at indicated intensities (black circles: medians, n = 7, Friedman test, **p < 0.01, ***p < 0.001). **(B)** Quantification of the rheobase shift (left) and the relative change in the number of ramp-evoked action potentials (right) in BiPOLES expressing cells. The injected current at the time of the first action potential was defined as the rheobase. Illumination with 490 nm light of increasing intensities activated somBiPOLES and BiPOLES-mediated $Cl^-$ currents shifting the rheobase to higher values and shunting action potentials (black circles: medians, n = 6, Friedman test, *p < 0.05, **p < 0.01, ***p < 0.001).

**Fig. S6**

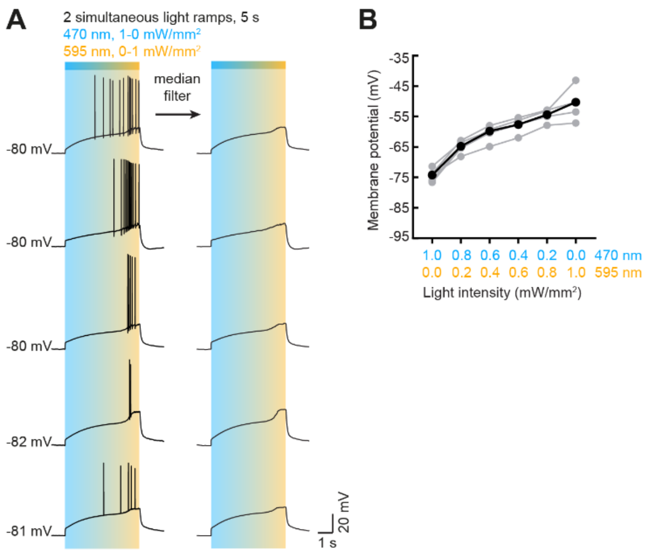


**Figure S6: Optical control of the neuronal membrane potential with somBiPOLES. (A)** Representative membrane voltage traces from five somBiPOLES-expressing neurons. In IC experiments, 470 and 595 nm light ramps of opposite gradient were applied as indicated. Initial high blue light caused a small depolarization, which increased steadily with an increasing 595/470 nm ratio eventually passing the action potential threshold, leading to action potential firing. Voltage traces were median-filtered to reveal the slow change in membrane voltage during the ramp protocol. **(B)** Quantification of membrane voltage at different 595/470 nm light intensity ratios. Despite different action potential thresholds, membrane potentials were highly similar between neurons at given light ratios, indicating that somBiPOLES is suitable to clamp the membrane voltage to a desired value by choosing the appropriate activation ratio of a cation (Chrimson) and anion (*Gt*ACR2) conductance (black circles = medians, n = 5).

**Fig. S7**

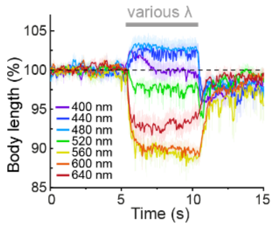


**Figure S7: Precise timing of bidirectional control of cholinergic motor neurons in *C. elegans*. (A)** Temporal dynamics of relative changes in body length upon illumination with light at wavelengths ranging from 400 to 640 nm in *C. elegans* expressing BiPOLES in cholinergic motor neurons (1.1 $mW/mm^2$, n = 9 - 14).

Fig. S8

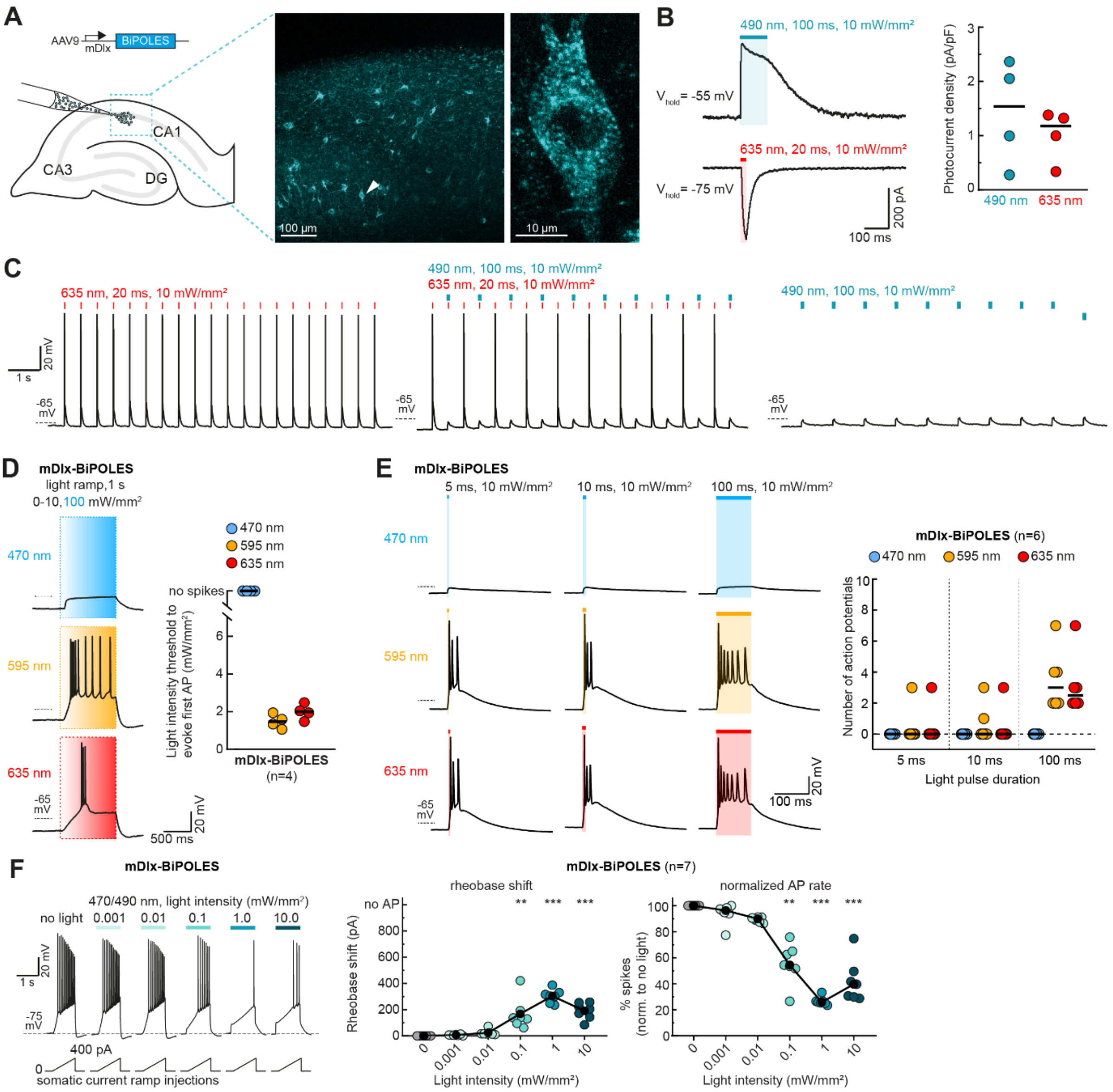


**Figure S8: Virally expressed mDlx-BiPOLES enables bidirectional control of GABAergic neuronal activity. (A)** Viral transduction of mDlx-BiPOLES in hippocampal organotypic slice cultures. Right: Maximum-intensity projection images of 2-photon stacks showing expression of BiPOLES in GABAergic neurons in CA1. Magnified view of a single neuron indicated by white arrowhead is shown on the right. **(B)** Left: Representative photocurrent traces measured in an mDlx-BiPOLES-expressing CA1 GABAergic neuron. Photocurrents evoked by a 490 nm light pulse (100 ms, 10 mW/mm$^2$) were recorded at a membrane voltage of -55 mV and photocurrents evoked by a 635 nm light pulse (20 ms, 10 mW/mm$^2$) were recorded at a membrane voltage of -75 mV. Right: Quantification of photocurrent densities evoked under the indicated conditions (black horizontal lines: medians, n = 4). **(C)** IC characterization of bidirectional optical spiking-control with mDlx-BiPOLES. Voltage traces showing red-light-evoked action potentials (left), which were blocked by a concomitant blue-light pulse (middle). Blue light alone did not trigger action potentials (right). Small depolarizations seen with blue light are due to the slightly depolarized chloride Nernst potential in our recording conditions. **(D)** Left: Representative IC membrane voltage traces measured in mDlx-BiPOLES-expressing neurons. In IC experiments, light ramps were applied as indicated. The intensity was ramped linearly over 1 s from 0 to 10 mW/mm$^2$ or to 100 mW/mm$^2$ for 470 nm to rule out the possibility that high-intensity blue light might still evoke action potentials. Right: Quantification of the light intensity threshold at which the first action potential (AP) was evoked. 470-nm light up to 100 mW/mm$^2$ did not evoke action potentials in mDlx-BiPOLES-expressing cells, while 595 and 635 nm light evoked action potentials at intensities comparable to pyramidal cells expressing BiPOLES. **(E)** Extended duration of illumination increases the probability and number of action potentials. Left: Representative IC membrane voltage traces measured in mDlx-BiPOLES-expressing neurons illuminated as indicated. Right: quantification of the number of action potentials evoked by the different illumination protocols (black horizontal lines: medians, n = 6). **(F)** IC characterization of mDlx-BiPOLES-mediated neuronal silencing. Current ramps (from 0–100 to 0–900 pA) were injected into mDlx-BiPOLES-expressing cells to induce action potentials. The injected current at the time of the first action potential was defined as the rheobase. Illumination with blue light of increasing intensities (from 0.001 to 10.0 mW/mm$^2$) activated BiPOLES-mediated $Cl^-$ currents shifting the rheobase to higher values. Middle: Quantification of the rheobase shift at different light intensities. Right: Relative change in the number of ramp-evoked action potentials upon illumination with blue light at indicated intensities (black circles: medians, n = 7, Friedman test, **p < 0.01, ***p < 0.001).

**Fig. S9**

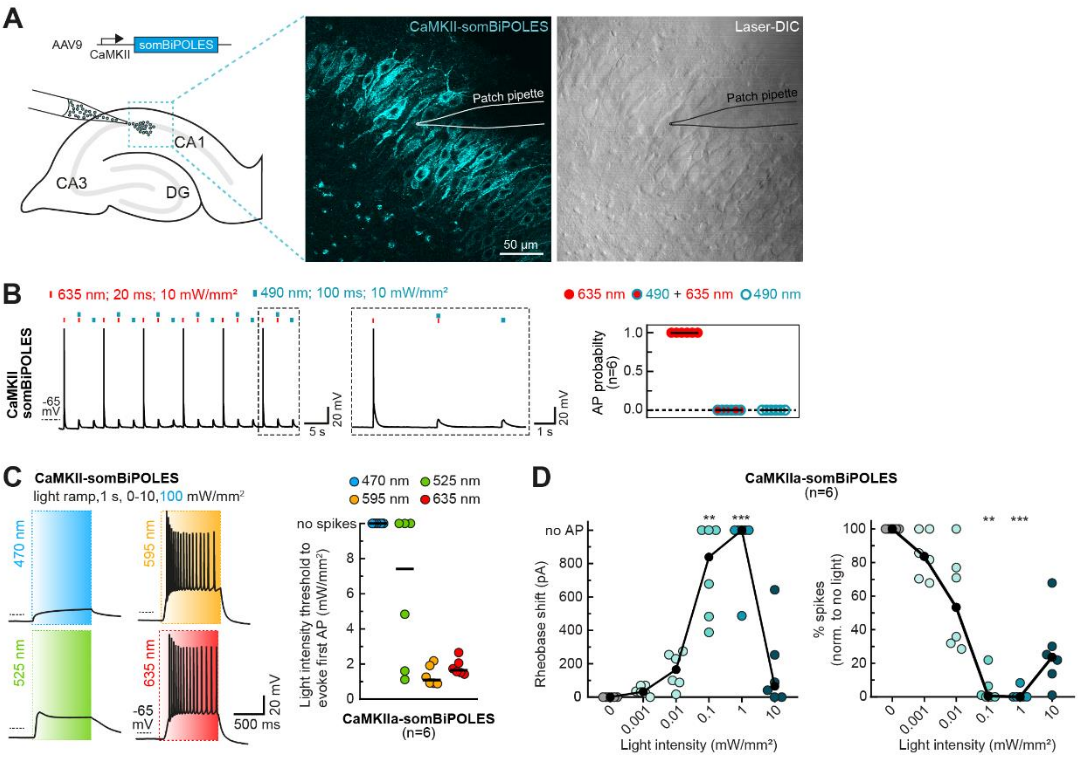


**Figure S9: Virally expressed CaMKII-somBiPOLES enables bidirectional control of activity in projection neurons. (A)** Viral transduction of CaMKII-somBiPOLES in hippocampal organotypic slice cultures. Right: Single-plane 2-photon fluorescence (cyan) and laser-DIC (gray) images showing expression of somBiPOLES in pyramidal cells of *stratum pyramidale* and cellular morphology, respectively. The position of the patch pipette is depicted by a drawing of its outline. **(B)** Current-clamp (IC) characterization of bidirectional optical spiking-control with CaMKII-somBiPOLES. Left: Voltage traces showing red-light-evoked action potentials, which were blocked by a concomitant blue-light pulse. Blue light alone did not trigger action potentials. Right: quantification of action potential probability under indicated conditions (black horizontal lines: medians, n = 6). **(C)** Left: Representative membrane voltage traces measured in CaMKII-somBiPOLES-expressing pyramidal neurons. In IC experiments, light ramps were applied as indicated. The intensity was ramped linearly from 0 to 10 mW/mm$^2$ over 1 s, except for 470 nm ramps, which were ranging to 100 mW/mm$^2$ to rule out the possibility that high-intensity blue light might still evoke action potentials. Right: Quantification of the light intensity threshold at which the first action potential was evoked (black horizontal lines: medians, n = 6). **(D)** IC characterization of CaMKII-somBiPOLES-mediated neuronal silencing. Current ramps (from 0–100 to 0–900 pA) were injected into CaMKII-somBiPOLES-expressing cells to induce action potentials. The injected current at the time of the first action potential was defined as the rheobase. Illumination with blue light of increasing intensities (from 0.001 to 10.0 mW/mm$^2$) activated somBiPOLES-mediated $Cl^-$ currents shifting the rheobase to higher values. Note the apparent absence of a rheobase shift at 10 mW/mm$^2$, which is likely due to the added depolarization mediated by Chrimson at high light intensities. Nonetheless, following action potentials with increasing current injection, as observed in absence of blue light, were efficiently shunted (black circles: medians, n = 6, Friedman test, $^{**}p < 0.01$, $^{***}p < 0.001$).

**Fig. S10**

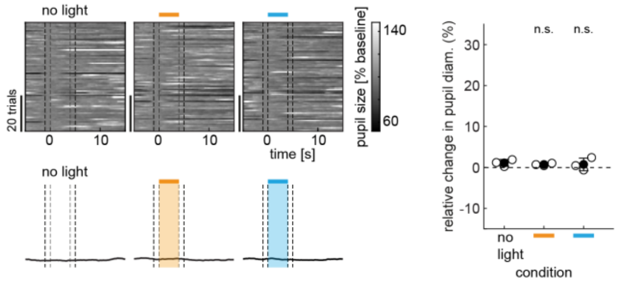


**Figure S10: Pupil dilation is not altered by light applied to the LC in wild-type animals.** Quantification of normalized pupil size in one wild-type animal under various stimulation conditions as indicated. Orange and blue bars indicate time of illumination with 594 (orange) and 473 nm (blue), respectively. Top left: single trials. Bottom left: mean ± SEM. Dashed lines show time points used for quantification in the plot on the right. Right: quantification of relative pupil size (n = 3 mice, n.s. = not significant).
